# Ethanol stimulates trehalose production through a SpoT-DksA-AlgU dependent pathway in *Pseudomonas aeruginosa*

**DOI:** 10.1101/523126

**Authors:** Colleen E. Harty, Dorival Martins, Georgia Doing, Dallas L. Mould, Michelle E. Clay, Dao Nguyen, Deborah A. Hogan

**Affiliations:** Microbiology and Immunology, Geisel School of Medicine at Dartmouth, Hanover, NH, USA.; Meakins-Christie Laboratories and Translational Research in Respiratory Diseases Program, Research Institute of the McGill University Health Centre, Montréal, QC H4A 3J1, Canada; Department of Medicine, McGill University, Montréal, QC H4A 3J1, Canada

**Author notes:** To whom correspondence should be addressed, Department of Microbiology and Immunology, Geisel School of Medicine at Dartmouth Rm 208 Vail Building, Hanover, NH 03755.

## Abstract

*Pseudomonas aeruginosa* frequently resides among ethanol-producing microbes, making its response to these microbially-produced concentrations of ethanol relevant to understanding its biology. Our ranscriptome analysis found that the genes involved in trehalose metabolism were induced by low concentrations of ethanol, and levels of intracellular trehalose increased significantly upon growth with ethanol. The increase in trehalose was dependent on the TreYZ pathway, but not other trehalose metabolic enzymes TreS or TreA. The sigma factor AlgU (AlgT), a homolog of RpoE in other species, was required for increased expression of the *treZ* gene and trehalose levels, but induction was not controlled by the well-characterized proteolysis of its antisigma factor MucA. Growth with ethanol led to increased SpoT-dependent (p)ppGpp accumulation, which stimulates AlgU-dependent transcription of *treZ* and other AlgU-regulated genes through DksA, a (p)ppGpp and RNA polymerase binding protein. Ethanol stimulation of trehalose also required acylhomoserine lactone (AHL)-mediated quorum sensing, as induction was not observed in a Δ*lasR*Δ*rhlR* strain. A network analysis using a model, eADAGE, built from publicly available *P. aeruginosa* transcriptome datasets (1) provided strong support for our model that *treZ* and co-regulated genes are controlled by both AlgU and AHL-mediated QS (QS). Consistent with (p)ppGpp and AHL-mediated quorum sensing regulation, ethanol, even when added at the time of culture inoculation, stimulated *treZ* transcript levels and trehalose production in cells from post-exponential phase cultures but not from exponential phase cultures. These data highlight the integration of growth and cell density cues in the *P. aeruginosa* transcriptional response to ethanol.

**Importance:** *Pseudomonas aeruginosa* is often found with bacteria and fungi that produce fermentation products including ethanol. At concentrations similar to those produced by environmental microbes, we found that ethanol stimulated expression of trehalose biosynthetic genes and cellular levels of trehalose, a disaccharide that protects against environmental stresses. The induction of trehalose by ethanol required the alternative sigma factor AlgU through DksA and SpoT-dependent (p)ppGpp. Trehalose accumulation also required AHL quorum sensing and only occurred in post-exponential phase cultures. This work highlights how cells integrate cell-density and growth cues in their responses to products made by other microbes and a reveals a new role for (p)ppGpp in the regulation of AlgU activity.

## Introduction

*Pseudomonas aeruginosa* is a ubiquitous Gram-negative bacterium that can cause acute and chronic infections in a broad range of hosts. *P. aeruginosa* frequently causes chronic infections in individuals with the genetic disorder cystic fibrosis (CF). Diverse bacterial and fungal taxa often co-infect with *P. aeruginosa* in CF airways (2-6), and many of these taxa are robust fermenters capable of ethanol production (7, 8). Ethanol has also been identified as a volatile biomarker in exhaled breath condensates that discriminates between healthy individuals and those with CF (9).

Ethanol has a range of biological activities that vary based on its concentration. At concentrations in the 3-5% range and higher, ethanol can inhibit growth or kill *P. aeruginosa* (10-12). The effects of biologically-produced concentrations of ethanol within the range experienced by organisms in polymicrobial communities (0.1-1.1%) (13-21) have been less well studied. Several studies have shown that 1% ethanol can alter pathogenesis and interspecies interactions (13, 15, 17, 19). In *Acinetobacter baumannii*, ethanol enhances virulence toward *Caenorhabditis elegans* (17) and *Galleria mellonella* (19). This may be due to enhanced production of cytotoxic phospholipase C and increased expression of nutrient uptake pathways (18). In *P. aeruginosa*, ethanol produced by *C. albicans* influences the expression of the antifungal phenazine 5-methyl phenazine-1-carboxylic acid (5MPCA) and 1% ethanol was sufficient to modulate phenazine production and stimulate the exopolysaccharide Pel and Psl-mediated biofilm and pellicle formation (13, 16).

The *P. aeruginosa* alternative sigma factor, AlgU, also named AlgT, has been well-studied for its positive regulation of production of alginate, an exopolysaccharide (22, 23). AlgU is an extracytoplasmic sigma factor that is homologous to RpoE (σ^E^ or σ^22^) in other Gram-negative bacteria (24). *P. aeruginosa* mutants lacking *algU* have increased resistance to hydrogen peroxide compared to alginate over-producing mucoid counterparts due to transcriptional de-repression of catalase *katA*, but are more susceptible to host antimicrobial peptides (25, 26). In other species, σ^E^ is necessary for fitness in response to high concentrations of ethanol (3-10%) and inhibitory concentrations of salt (27-30). A well-known mechanism of activation of σ^E^ in response to stresses that perturb the cell envelope is by proteolytic degradation of its anti-sigma factor by specific proteases (31, 32). In *P. aeruginosa*, the AlgU anti-sigma factor is MucA, and *mucA* mutations lead to high AlgU activity. Naturally-occurring *mucA* mutants are frequently observed in populations from chronic *P. aeruginosa* lung infections and strains overproduce alginate (33, 34).

In *E. coli* and *Salmonella enterica*, σ^E^ activity can also be modulated by the alarmone (p)ppGpp (35, 36). (p)ppGpp is an intracellular molecular signal that is synthesized by either the synthase RelA or a hybrid synthase/hydrolase, SpoT (37) in response to nutrient limitation and various environmental stressors (38). (p)ppGpp can complex with the RNA polymerase binding protein DksA to promote transcription initiation and elongation and alter the effects of RNA polymerase-associated sigma factors including RpoE (39, 40).

In this work, we show that a sub inhibitory concentration of ethanol (1%) induces the expression of genes involved in the metabolism of trehalose and biochemical assays found significant increases in intracellular trehalose, a disaccharide that serves as both a compatible solute and a carbon source. Increased trehalose in response to ethanol required the TreYZ trehalose biosynthetic enzymes, but not the TreS trehalose synthase. Ethanol induction of *treZ* gene expression and trehalose requires the sigma factor AlgU. AlgU was not activated by release from MucA, but rather in a manner dependent on (p)ppGpp. Ethanol caused a 2.5-fold increase in (p)ppGpp levels, which was specifically dependent on SpoT (p)ppGpp synthase, and the (p)ppGpp-binding protein DksA was required for ethanol-induced stimulation of *treZ* gene expression and trehalose levels. Consistent with previous reports (41, 42), acylhomoserine lactone-mediated (AHL) quorum sensing was also required for transcriptional induction of the *treZ* gene by ethanol, as a Δ*lasR*Δ*rhlR* mutant defective in AHL-mediated quorum sensing did not show increased trehalose levels in response to ethanol. The stimulation of trehalose levels by salt did require AlgU, and trehalose stimulation in response to salt was lower, but still occurred, in mutants lacking the factors necessary for the response to ethanol. Analysis of genes differentially expressed when ethanol is in the growth medium, performed using the eADAGE gene expression model constructed with data from over 1056 different samples (1), placed the *treYZ* genes among a cluster of co-regulated genes within the AHL-controlled quorum sensing (QS) and AlgU regulons. Ethanol, even when added at the time of culture inoculation, only stimulated AlgU-regulated genes and trehalose production in cells during post-exponential phase, which is consistent with our model that regulators that monitor growth and cell density cues are integrated into the *P. aeruginosa* response to ethanol.

## Materials and Methods

### Strains and growth conditions

Bacterial strains and plasmids used in this study are listed in Table S1. Bacteria were maintained on 1.5% agar LB (lysogeny broth) plates (43). Where stated, ethanol (200-proof) was added to the medium (liquid or molten agar) to a final concentration of 1% unless otherwise stated. NaCl to a final concentration of 500 mM was added to liquid medium as indicated. Mutants from the PA14 Non-Redundant (NR) Library were grown on LB with 60 μg/mL gentamicin (44). When strains from the NR library were used, the location of the transposon insertion was confirmed using site-specific primers. The primers are listed in Table S2. Where stated, LB medium was buffered to pH 8 with 100 mM HEPES buffer (referred to as buffered LB). M63 medium contained 0.2% glucose and 2% casamino acids (45). When ethanol was supplied as a sole carbon source, glucose and amino acids were omitted.

Planktonic cultures were grown at 37°C on a roller drum.

### Construction of in-frame deletions, complementation, and plasmids

Construction of plasmids, including in-frame deletion and complementation constructs, was completed using yeast cloning techniques in *Saccharomyces cerevisiae* as previously described (46) unless otherwise stated. Primers used for plasmid construction are listed in Table S2. In-frame deletion and single copy complementation constructs were made using the allelic replacement vector pMQ30 (46). Deletions of *relA* and *spoT* were introduced in the PA14 strain using the pEX18Gm suicide vector to create unmarked deletion mutants (47), as previously described (48). Promoter fusion constructs were made using a modified pMQ30 vector with *lacZ*-*GFP* fusion integrating at the neutral *att* site on the chromosome.

The *rhlI* promoter region was amplified from PA14 gDNA using the Phusion High-Fidelity DNA polymerase with primer tails homologous to the modified pMQ30 ATT KI vector containing the *lacZ-gfp* reporters. The 195 bp upstream promoter region includes a RhlR-binding site as annotated by pseudomonas.com at positions –192 to +3. All plasmids were purified from yeast using Zymoprep™ Yeast Plasmid Miniprep II according to manufacturer’s protocol and transformed into electrocompetent *E. coli* strain S17 by electroporation. Plasmids were introduced into *P. aeruginosa* by conjugation and recombinants were obtained using sucrose counter-selection and genotype screening by PCR.

### *P. aeruginosa* growth assays

For growth curves shown in Figure S1, overnight cultures were diluted into 5 mL fresh buffered LB medium in 18 × 150 mm borosilicate glass tubes without or with 1% ethanol to an OD_600nm_ of ∼0.05 and incubated at 37°C on a roller drum. Similar culture vessels and volumes were used in all assays unless otherwise specified. OD_600_ measurements were taken on a Genesys 6 spectrophotometer. For data in Figure S4, overnight cultures were diluted into fresh LB medium buffered to pH 8 with HEPES without or with 1% ethanol to an OD_600_ of ∼0.05 and 150 µl of the cell suspension was pipetted into 96-well plates. Plates were grown at 37°C with continuous shaking at ∼150 rpm. OD_600_ measurements were taken at 16 h of incubation using a microplate reader.

### Microarray analysis

Cultures of *P. aeruginosa* PA14 wild type were grown overnight in LB at 37°C on a roller drum. 5 μL of overnight culture were spotted onto T-broth plates (1.5% agar) (49) ± 1% ethanol. Plates were incubated at 37° for 16 h. Colonies were scraped up from the plates for RNA isolation using the Qiagen RNeasy Mini kit. The samples were DNase treated using the Invitrogen Turbo DNA-Free kit. As previously described (50), cDNAs labeled with biotin-ddUTP (Enzo Bio-Array terminal labeling kit, Affymetrix) were hybridized to Pseudomonas GeneChips using the GeneChip fluidics station 450 (Affymetrix) according to manufacturer’s instructions. GeneChips were scanned in the Dartmouth Genomics and Microarray Laboratory using the GeneChip Scanner 3000 7G (Affymetrix) and the BioConductor Affy library was used to read CEL file data. Data were normalized with RMA in BioConductor (51).

### eADAGE analysis

Genes upregulated 2-fold or more were analyzed within the context of their expression patterns in a compendium of 1051 publicly available microarrays from the Gene Expression Omnibus as determined by a machine learning model, eADAGE (1). Pearson correlations greater than 0.5 between the learnt parameters corresponding to each gene were visualized by edge weights in the resultant network (52). Genes differentially expressed between a wild type and Δ*lasR*Δ*rhlR* mutant strain sampled at different time points over the course of growth are referred to as the quorum sensing (QS)-controlled regulon (41) (Table S3D). The QS-controlled genes are presented as a network, plotted using the Fruchterman-Reingold force-directed algorithm, in which correlations in gene expression were indicated with the presence of edges and genes with shorter edges are more strongly correlated in expression pattern. The network was generated in R using the “network” (53, 54), “GGally” (55) and “ggplot2” (56) packages. The genes within the QS-controlled gene set that were also differentially expressed upon deletion of *algU* (57) were indicated as green nodes and genes within the QS-controlled gene set that were also differentially expressed upon deletion of *rpoS* (42) (Table S3E) are indicated as pink nodes. QS-controlled genes that were not also differentially expressed upon deletion of AlgU or RpoS are presented as blue nodes. The complete gene lists for each data set and accompanying R code are available as a supplemental file. If necessary, PA14 gene numbers gene were converted to PAO1 ortholog gene numbers, and PAO1 gene numbers were converted to gene names, using *Pseudomonas aeruginosa* PA14 109 orthologs and *Pseudomonas aeruginosa* PAO1 107 annotations from www.pseudomonas.com (58).

### Quantitative PCR analysis of transcripts

For quantitative real-time PCR experiments, cultures of indicated strains of *P. aeruginosa* were grown for 16 h in 5 mL of buffered LB at 37°C on a roller drum. RNA was isolated from planktonic cultures using the Qiagen RNeasy Mini kit. The samples were DNase-treated using the Invitrogen Turbo DNA-Free kit. cDNA was synthesized using the RevertAid H-minus first-strand synthesis kit using the GC-rich protocol with the following temperatures: 25°C for 5 minutes, 50°C for 60 minutes, and 70°C for 5 minutes. Synthesized cDNA was diluted 1:5 in molecular grade water and stored at −20°C. Quantitative PCR expression analysis was performed using an Applied Biosystems 7500 Real-Time PCR system with BioRad SsoFast Evagreen Supermix and primers listed in Table S2. A cycling regimen of 95°C for 30 seconds; 39 cycles of 95°C for 10 seconds and 60°C for 5 seconds; and a final 65°C for 3 seconds was used. Experimental transcripts were normalized to the housekeeping gene *rpoD*.

### Measurement of trehalose in cells

Trehalose was quantified from whole cell lysates as described previously (59, 60) with slight modifications. Briefly, bacterial cultures were grown for 16 h in LB or M63 medium without or with 1% ethanol as stated, in 18mm borosilicate culture tubes at 37°C on a roller drum. Cultures were inoculated from strains grown on LB plates. A 250 μL volume of culture was concentrated to an OD_600_ of 8.0 in sterile water. Cell suspensions were boiled for 10 minutes to lyse cells. The resulting lysate was centrifuged at 16,000 xg, and 100 μL aliquots of lysate were transferred to new tubes. One tube of lysate was treated with 1 μL of trehalase (Sigma-Aldrich) enzyme or vehicle control. Glucose in the lysate samples was quantified using the glucose oxidase kit (Sigma; catalog no. GAGO20). Trehalose concentrations were calculated based on a standard curve and subtraction of the basal glucose and expressed relative to OD units.

### (p)ppGpp measurments

Cultures inoculated at OD_600_= 0.05 were grown for 16 h to an OD_600_∼2.0 in 5 mL M63 medium without and with 1% ethanol in 18 mm culture tubes at 37°C with 250 rpm shaking. About 2 mL of cultures were pelleted at 10,000xg for 5 min and their (p)ppGpp was extracted as described previously (61). Briefly, the pellets were suspended in 200 µL of 10 mM Tris-HCl pH 7.8 containing 1 mg/mL lysozyme and 15 mM magnesium acetate. The suspensions were vortexed for 3 s and subjected to two freeze-thawing cycles. Then, 15 µL of a 10% deoxycholate solution was added and the suspensions were vortexed for 15 s. Finally, 200 µL of 10 mM Tris-HCl pH 7.8 containing 15 mM magnesium acetate was added and the samples were centrifuged at 12,000xg for 10 min. The supernatants were used for (p)ppGpp quantification, which was performed using the chemosensor Bis-Zn^2+^-dipicolylamine (PyDPA) as previously described (62). The measurements of (p)ppGpp-PyDPA complex were carried out immediately after the probe addition using fluorescence spectroscopy (Ex/Em= 344/480 nm) in a Tecan Infinite M1000 plate reader. To account for the interference of other nucleotides, Δ*relA*Δ*spoT* mutant extracts were used and the absolute (p)ppGpp values were determined using a calibration curve with purified (p)ppGpp (TriLink Biotechnologies) spanning a linear range of 0.4-6 µM (p)ppGpp. For normalized (p)ppGpp measurements, fluorescence at Ex/Em 344/480 nm was normalized by the fluorescence at Ex/Em 344/380 nm (free nucleotides complex with PyDA) and by the OD_600_ of the washed cultures.

### Measurement of β-galactosidase in reporter fusion strains

Cells with an *rhlI* promoter fusion to *lacZ-GFP* genes integrated at the *attB* locus were grown in 2 mL of LB overnight culture grown at 37°C were pelleted, washed twice and re-suspended in PBS. The washed cells were diluted to a starting OD_600_=0.05 in 5 mL of M63 medium without and with 1% EtOH. After 16 h on a roller drum at 37°C, β-Galactosidase (β-Gal) activity was measured as described by Miller (63).

### Statistics

Unless otherwise stated, data are based on three biological replicates with the mean and standard deviations calculated, and are representative of at least three independent experiments containing multiple replicates. Unless stated otherwise, means and standard deviations were calculated in Graph Pad Prism 8 and analyses were completed using a two-way ANOVA and Tukey’s multiple comparisons test, with p-values indicated in figure legends. Regulon enrichments were determined by hypergeometric tests using the “phyper” function in the “stats” package in R.

### Accession Number

Data for our microarray analysis of *P. aeruginosa* PA14 wild type in response to ethanol has been uploaded to the GEO repository (https://www.ncbi.nlm.nih.gov/geo/) with the accession number GSE124852.

## Results

### Analysis of the transcriptome upon growth in the presence of ethanol

To examine the transcriptional response of *P. aeruginosa* to 1% ethanol, RNA from *P. aeruginosa* grown as colony biofilms for 16 h on tryptone agar ±1% ethanol was analysed using *P. aeruginosa* Affymetrix GeneChips. Similar to published results (13), the presence of 1% ethanol in the medium did not affect the number of CFUs in colony biofilms (Fig. S1A).

Fifty-four transcripts were higher by two-fold or more in cells grown with ethanol, with an FDR-corrected p-value less than 0.05, and twenty genes were found to be lower by 2-fold or more in the presence of ethanol (Table S3A and B). Among the most differentially expressed genes were those involved in trehalose metabolism (*treZ* (3-fold), *treA* (2.1-fold) and *treS* (2.5-fold) (64). Other genes that were changed upon growth with ethanol are discussed in more detail below. To determine if ethanol also led to increased levels of trehalose, intracellular trehalose concentrations were measured in cells grown ± ethanol in LB, a nutrient-rich medium, and M63, a defined medium. The LB was pH-buffered with HEPES because we observed that ethanol led to a lower final pH in *P. aeruginosa* cultures grown with ethanol (final pH of 8.3 in control and pH 6.5 in ethanol from an initial pH of 7.1), despite similar growth kinetics (Fig. S1B), as has been described in *E. coli* (65) and *Acinetobacter baumannii* (19). Ethanol led to significant increases in trehalose in both buffered LB (>2-fold higher with ethanol) and in M63 (20-fold higher with ethanol).

### Increased trehalose in response to ethanol requires *treYZ* genes

In Pseudomonads such as *P. aeruginosa* and *Pseudomonas syringae*, trehalose can be synthesized by the TreYZ pathway, which converts glycogen to trehalose via a maltooligosyl trehalose synthase and glycosyl hydrolase (66), and by TreS, a trehalose synthetase that uses maltose as a substrate (Fig. 1A for schematic) (64, 67). Trehalose is degraded by the trehalase TreA; in other species, TreS also has trehalose catabolic activities (68, 69).

**Figure 1.**
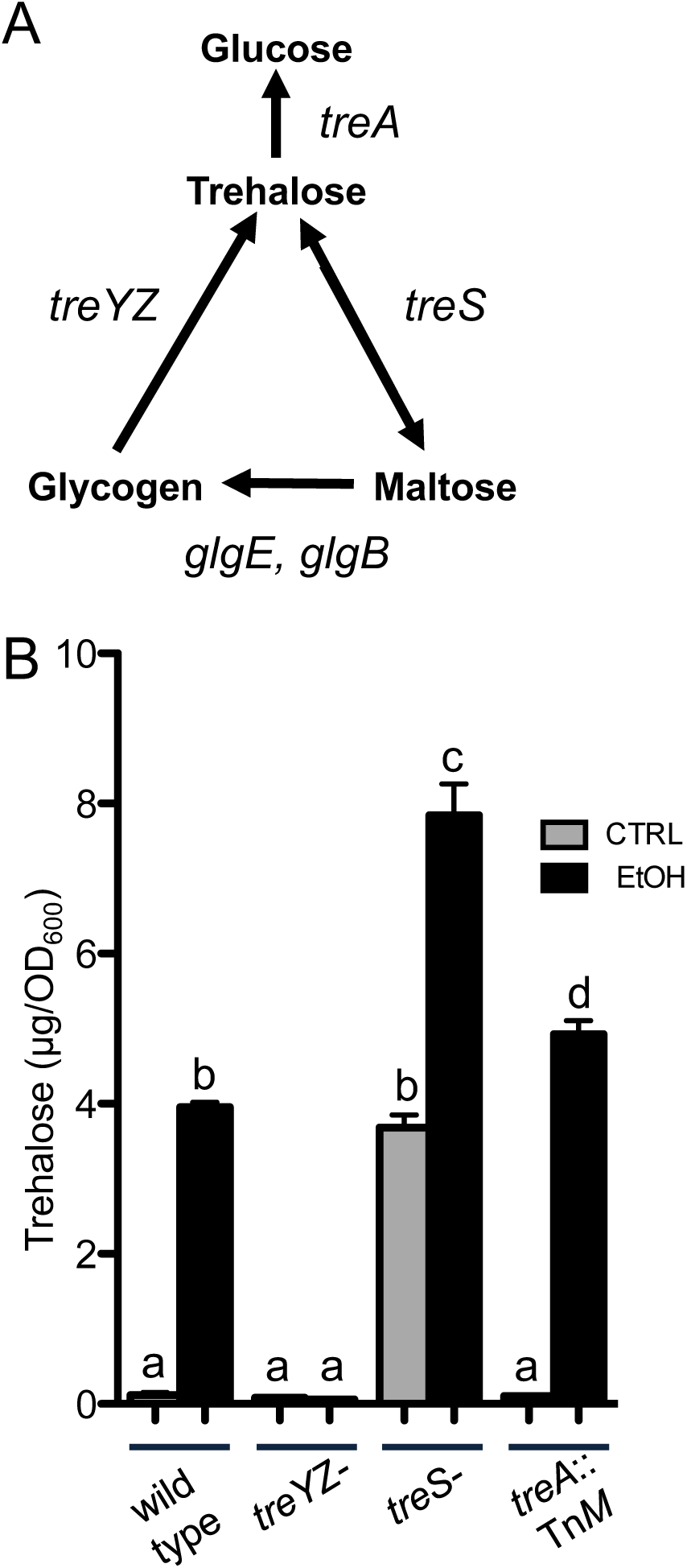
Trehalose accumulation in response to ethanol requires *treYZ*. A. Schematic of trehalose biosynthetic pathways in *P. aeruginosa.* B. Trehalose levels in trehalose metabolic mutants grown in M63 medium without and with 1% ethanol for 16 hours. Data are representative of at least 3 independent experiments with 3 biological replicates each. a-b, a-c, a-d, b-c, c-d p<0.0001; b-d p<0.02 based on two-way ANOVA and Tukey’s multiple comparisons test.

To determine which metabolic pathway was responsible for increased trehalose in cells grown with ethanol, we used mutants lacking the 6-gene operon (PA14_36570-PA14_36630) that contains *treY* and *treZ* genes (referred to here as *treYZ*^*-*^ (64)), the 3-gene operon that contains *treS*^*-*^ (referred to here as *treS*^*-*^ (64)), and an insertional mutant of *treA* (44, 64). While *treS*^*-*^ *and treA*::Tn*M* both showed a marked increase in trehalose in the presence of ethanol in comparison to controls, the *treYZ*^*-*^ strain did not, suggesting the increase in trehalose was by the TreYZ pathway (Fig. 1B). The significantly higher levels of trehalose in the *treS*^*-*^ mutant in both control and ethanol conditions relative to the wild type suggested that the TreS catabolic activity was present under these conditions (70).

Ethanol catabolism occurs mainly by the pyrroloquinoline quinone (PQQ)-dependent alcohol dehydrogenase, ExaAB, and the resultant acetaldehyde is catabolized through a pathway that includes acetyl-CoA synthetase (AcsA). We have shown previously that ethanol catabolic mutants (*exaA, pqqB*, and *acsA*) are defective in growth on ethanol as a carbon source (13). Ethanol catabolic mutants still showed a stimulation of trehalose production with ethanol in the growth medium (Fig. S2A).

Because we previously showed that ethanol (1%) led to increased production of the Pel exopolysaccharide through the diguanylate cyclase WspR (13), we determined if changes in trehalose occurred in response to changes in Pel production. We found that both the *pelA* and *wspR* mutants still had higher levels of trehalose in cells grown with ethanol (Fig. S2B) suggesting that changes in exopolysaccharide biosynthesis did not cause the increase in trehalose.

### Ethanol induction of *treZ* gene expression and trehalose levels are dependent on AlgU

Several lines of evidence led us to hypothesize that the alternative sigma factor AlgU controlled the induction of trehalose metabolic genes in response to ethanol. First, *treZ, treS,* and *treA* have been reported to be differentially expressed in *algU* mutant strains when compared to a wild-type strain in transcriptomics studies (57, 71). Second, *osmC* and *pfpI*, two well-characterized members of the AlgU regulon (71-73) were differentially expressed in our transcriptomics analysis of cells grown ±1% ethanol (Table S3A).

In wild-type cells, qRT-PCR analysis found *treZ* to be 16-fold higher in cultures containing ethanol. In contrast, an in-frame Δ*algU* mutant had no significant difference in *treZ* expression between cells growth with and without ethanol (Fig. 2A). Like *treZ*, the expression of another AlgU regulated gene *osmC* was higher (8-fold) upon growth with ethanol and, as expected, the differential expression was dependent on AlgU (Fig. 2B). The Δ*algU* mutant also did not show an increase in trehalose upon growth with ethanol (Fig. 2C), and its defect could be complemented by restoring *algU* to the native locus (Fig. 2C). While the sigma factor AlgU was required for the induction of *treZ* transcripts and trehalose production, a mutant lacking another sigma factor, RpoS, which has been shown to regulate trehalose levels in *E. coli* (74), did not differ from the wild-type in its response to ethanol (Fig. S3).

**Figure 2.**
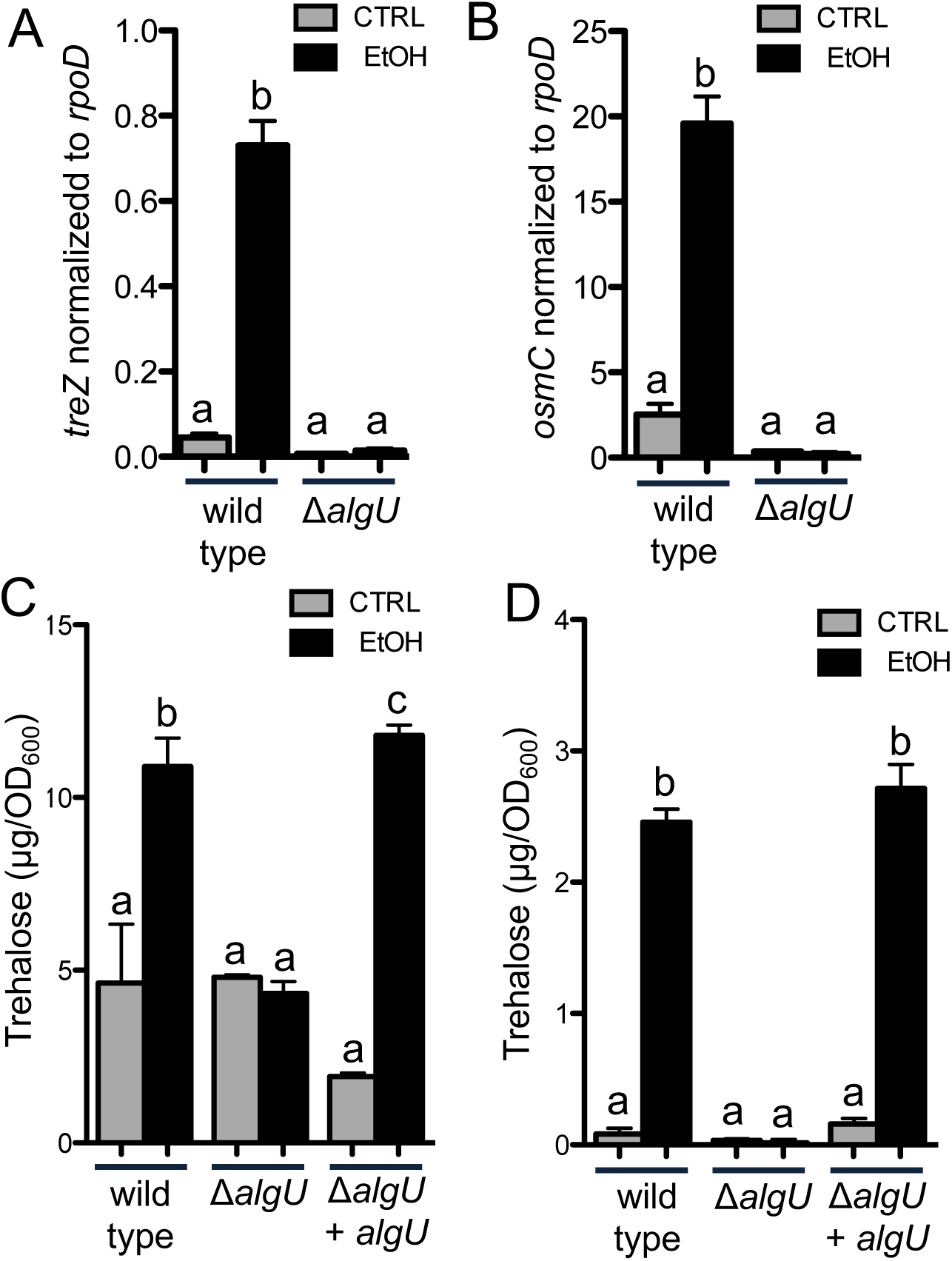
Analysis of the effects of ethanol on *treZ* and *osmC* transcript levels and intracellular trehalose in *P. aeruginosa* wild type and Δ*algU*. A and B. *treZ* (A) and *osmC* (B) transcript levels relative to *rpoD* after growth in the absence and presence of 1% ethanol in buffered LB for 16 hours a-b p<0.0001 two-way ANOVA and Tukey’s multiple comparisons test. C and D. Trehalose levels in wild type, δ*algU*, and δ*algU* + *algU* at the native locus in buffered LB (C) and M63 medium (D) without and with 1% ethanol for 16 hours. a-b p<0.002; a-c p<0.0009; b-c NS (C) a-b p<0.0001 (D) based on two-way ANOVA and Tukey’s multiple comparisons test. Data are representative of at least three independent experiments, each with at least three biological replicates.

### AlgU-dependent induction of trehalose in response to ethanol is independent of MucA cleavage and KinB regulation

AlgU activity is repressed by the anti-sigma factor MucA, and proteolysis of MucA is a well characterized means by which AlgU and its homologs are activated (25, 72, 73, 75-79). MucA is bound by the periplasmic protein MucB, which inhibits MucA cleavage (80-82). Several stimuli, including high concentrations of ethanol (>3%) (27, 28) can lead to periplasmic stress and RpoE activation by anti-sigma factor degradation in other species. Several lines of evidence suggest that MucA cleavage was not the mechanism by which ethanol stimulated levels of *treZ* mRNA and trehalose. First, we found that ethanol increased trehalose to a similar extent in wild type and *mucB* mutant cells in strain PA14 background (Fig. 3A); the *mucB* mutant has decreased stability of MucA and the strain is mucoid, which indicates higher AlgU activation of genes involved in alginate biosynthesis. Second, ethanol stimulated trehalose levels in *P. aeruginosa* strain PAO1 in which *mucA* contained the *mucA22* mutation, which is found frequently among clinical isolates to an extent similar to that observed in the wild type. The truncated MucA22 mutant, a variant frequently found among *P. aeruginosa* clinical isolates, is no longer regulated by proteolysis and no longer represses AlgU (83, 84). Lastly, ethanol also stimulated trehalose in the alginate-overproducing mucoid cystic fibrosis isolate FRD1 (85) which also has the *mucA22* allele (81, 86) (Fig. 3B). In FRD1, the stimulation of trehalose levels by ethanol required the presence of AlgU as the isogenic non-mucoid *algT/U*::Tn*501* (FRD440) derivative showed a large reduction in ethanol-stimulated trehalose (22) (Fig. 3C). These analyses showed that 1) ethanol induced responses similar to those in strain PA14 in genetically-distinct strains PAO1 and FRD1 and 2) that the increase in trehalose in response to ethanol is likely not due to the stimulation of MucA cleavage.

**Figure 3.**
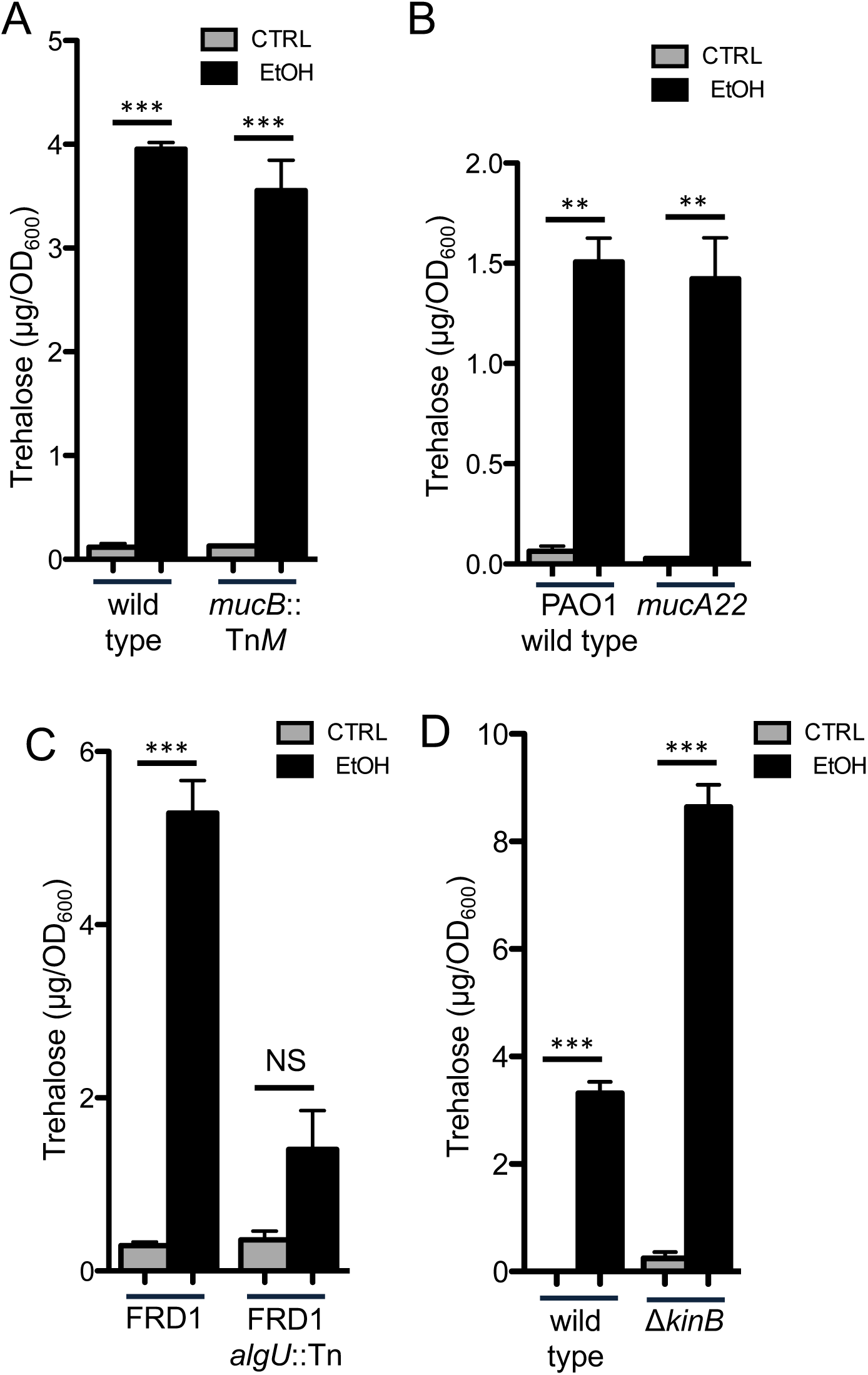
Trehalose levels in response to ethanol are independent of MucB, MucA cleavage, and KinB. Trehalose levels of A. *P. aeruginosa* strain PA14 wild type and the validated *mucB* transposon mutant. B. *P. aeruginosa* strain PAO1 wild type and a *mucA* (MucA22) mutant, C. FRD1 and its isogenic *algU::* Tn derivative. D. *P. aeruginosa* strain PA14 wild type and the Δ*kinB* mutant. Cultures grown in M63 medium without and with 1% ethanol for 16 hours. Data are representative of at least 2 experiments, each with 3 biological replicates. Statistics based on two-way ANOVA and Tukey’s multiple comparisons test *** p-value<0.0001; ** p≤0.0002.

A *kinB* loss-of-function has been associated with increased AlgU activity (72, 73) due to increased activity of the AlgB transcription factor that stimulates AlgU expression; KinB phosphatase activity normally represses AlgB (87). Thus, we determined if KinB was required for the difference in trehalose levels in cells grown with or without ethanol. While the *kinB* mutant consistently had higher levels of trehalose in control and ethanol conditions compared to the wild type, KinB was not required for the effects of ethanol on trehalose levels (Fig. 3D). Together, these data support our model that AlgU regulates trehalose through a mechanism that is not dependent on KinB.

### Ethanol stimulation of trehalose requires SpoT-generated ppGpp

In *E. coli,* the activity of RpoE, an AlgU homolog, is influenced by (p)ppGpp (35, 39, 40, 88, 89); (p)ppGpp effects on *P. aeruginosa* AlgU have not yet been reported. (p)ppGpp can be synthesized by either of two enzymes, RelA or SpoT (90-92). SpoT also has a (p)ppGpp degrading activity and, because very high levels of (p)ppGpp are toxic, mutants lacking *spoT* are only viable in the absence of RelA activity (93, 94). Thus, we determined if ethanol stimulation of trehalose levels was altered in either Δ*relA* or Δ*relA*Δ*spoT* strains. We found that while the Δ*relA* was like the wild type, ethanol did not influence trehalose levels in the Δ*relA*Δ*spoT* double mutant, suggesting that SpoT was required for trehalose accumulation in ethanol-grown cells (Fig. 4A). The induction of *treZ* and *osmC* were also dependent on SpoT (Fig. 4C and D).

**Figure 4.**
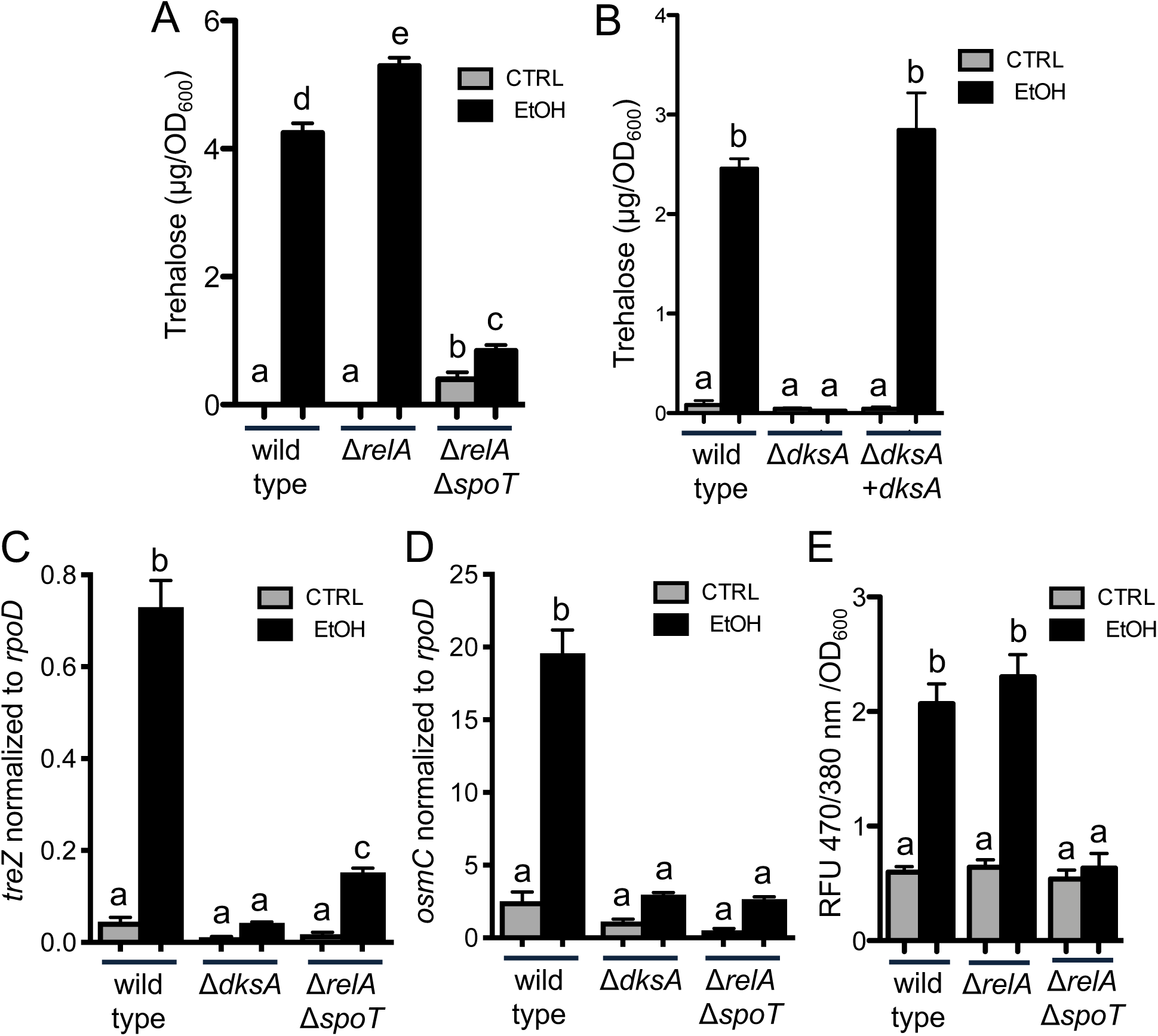
*dksA* and *spoT* are required for trehalose accumulation, *treZ* and *osmC* expression, and increased (p)ppGpp in response to ethanol. Trehalose levels of A. PA14 wild type, Δ*relA,* and Δ*relA*Δ*spoT* and B. PA14 wild type, Δ*dksA*, and Δ*dksA* + *dksA* in M63 medium without and with 1% ethanol for 16 hours. C. *treZ* and D. *osmC* transcripts normalized to *rpoD* in PA14 wild type, Δ*dksA*, and Δ*relAspoT* in buffered LB at 16 hours without and with 1% ethanol. A-D Data are representative of at least 3 independent experiments with at least 3 biological replicates each. E. (p)ppGpp quantification of PA14 WT, Δ*relA,* and Δ*relA*Δ*spoT* in M63 medium without and with 1% ethanol for 16 hours. Data are representative of at least 2 independent experiments. Statistics based on two-way ANOVA and Tukey’s multiple comparisons test. A. a-b p=0.0179; a-c, d-e p=0.0001; a-d, a-e, b-d, b-e, c-d, c-e p<0.0001; b-c = NS. B. a-b p<0.0001. C. a-b, b-c p<0.0001; a-c p≤0.05 D. a-b p<0.0001. E. a-b p<0.0001.

To determine if 1% ethanol altered (p)ppGpp levels in PA14 wild type, its levels were measured. We found that cells grown with ethanol had 3.45-fold higher levels of (p)ppGpp. The increase in (p)ppGpp was similar to the wild type in the *relA* mutant, but the Δ*relA*Δ*spoT* strain did not show an increase in (p)ppGpp in response to ethanol, indicating that ethanol stimulates (p)ppGpp production via SpoT (Fig. 4E).

### Ethanol stimulation of *treZ* and trehalose levels requires DksA

The (p)ppGpp signal influences the activity of RNA polymerase-sigma factor complexes through DksA, an RNA polymerase-binding protein. We found that a Δ*dksA* mutant no longer showed ethanol-induced trehalose accumulation and that the phenotype of the Δ*dksA* mutant was complemented by restoring *dksA* at the native locus (Fig. 4B). Consistent with trehalose measurements, ethanol-induced increases in *treZ* and *osmC* expression were greatly reduced in Δ*relA*Δ*spoT* and Δ*dksA* strains (Fig. 4C and D). Like the wild type, the growth kinetics of the Δ*dksA* mutant and Δ*relA*Δ*spoT* mutant were not reduced by 1% ethanol, though the Δ*dksA* mutant grew more slowly than the wild type in control conditions, consistent with published reports (95, 96) (Fig. S4).

### AlgU is required for, and SpoT and DksA contribute to, the induction of trehalose by high salt

In *P. aeruginosa* and other species, trehalose is induced by high salt (67, 97). Ausubel and colleagues (64) found that trehalose levels in high salt required TreYZ (64) and we confirmed these results (Fig. 5A and B). Furthermore, we found that trehalose induction in response to high salt was absent in the Δ*algU* mutant (Fig. 5C) as it was for ethanol (Fig. 1C and D). While (p)ppGpp and DksA were necessary for any induction of trehalose in response to ethanol, the Δ*dksA* and Δ*relAΔspoT* strains still showed a significant induction of trehalose in response to salt (Fig. 5C). The level of induction, however, was significantly lower in the Δ*dksA* and Δ*relAΔspoT* strains compared to wild type and Δ*relA* strain, suggesting that SpoT-dependent (p)ppGpp and the (p)ppGpp-responsive RNA polymerase binding protein DksA contributed to the response in cells from post-exponential phase cultures (Fig. 5C and D).

**Figure 5.**
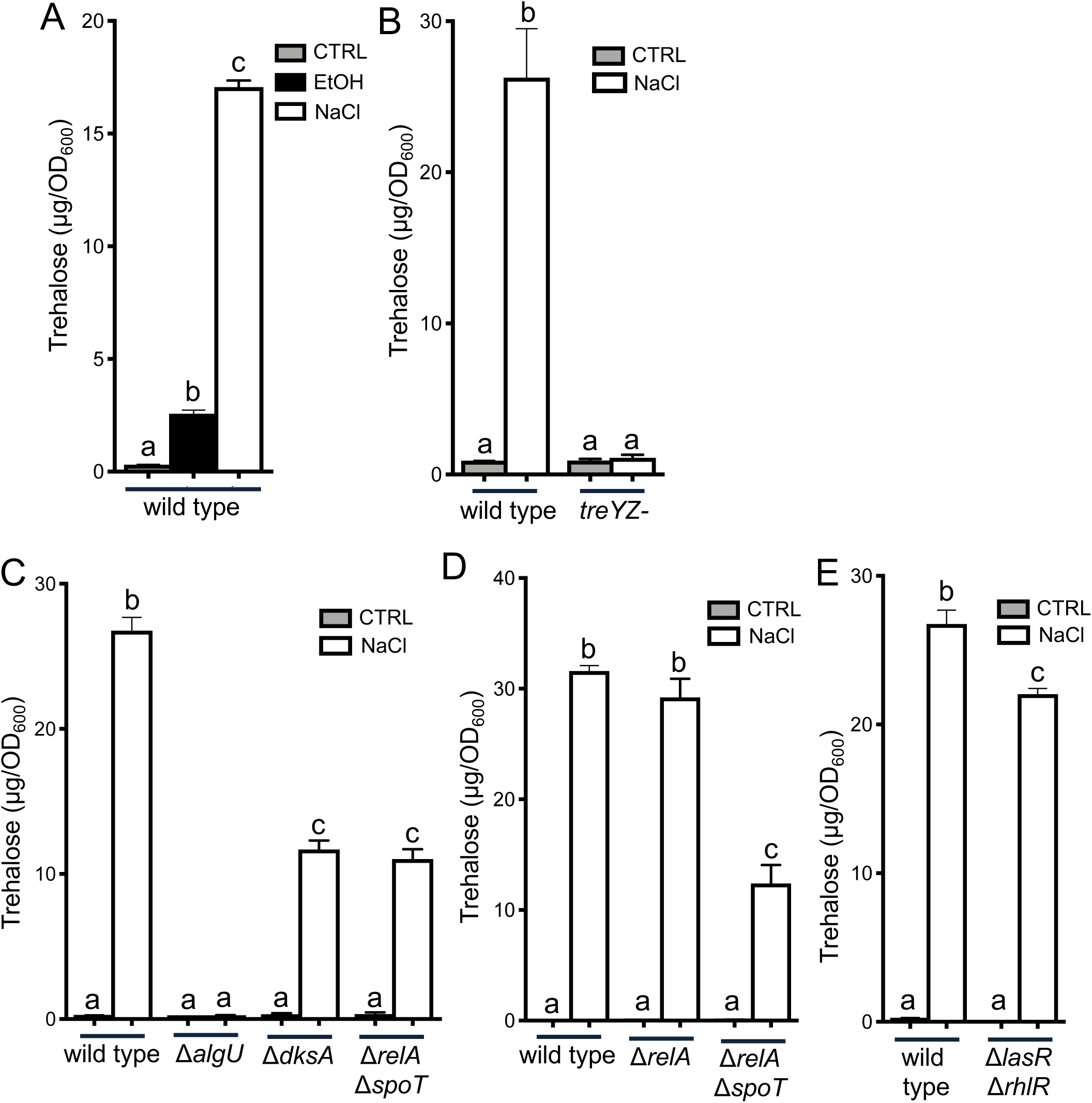
Trehalose accumulation in response to salt requires TreYZ and AlgU. A. 500 mM NaCl stimulates trehalose accumulation. B. Salt-stimulated accumulation of trehalose is dependent on the TreYZ pathway. C. Salt-stimulated trehalose accumulation requires *algU* and *dksA* and *spoT* contribute. D. Salt stimulated trehalose accumulation is independent of *relA*. E. Trehalose accumulation in salt is similar to wild type in Δ*lasR*Δ*rhlR*. Cultures were grown 16 hours in M63 medium with 1% ethanol or 500 mM NaCl as indicated. Data are representative of two or more experiments, each with at least 3 biological replicates. Statistics based on one-way ANOVA (A) or two-way ANOVA (B,C,D) and Tukey’s multiple comparisons test. A. a-b p=0.0005; a-c, b-c p<0.0001. B. a-b, p<0.0001. C. a-b, a-c, a-d, b-c, p<0.0001. D. a-b, a-c, a-d, b-c, p<0.0001. E. a-b, a-c p<0.0001; b-c p=0.0007

### Quorum sensing master regulators are necessary for the ethanol induction of trehalose

Schuster et al. (41, 42) found that *treZ* and other genes involved in trehalose biosynthesis and *osmC*, were at lower levels in a PAO1 Δ*lasR*Δ*rhlR* mutant compared to the wild type. They reported that *treZ* and *osmC* fell within a subset of QS-controlled genes that were induced later in growth in batch culture relative to other QS-controlled genes (41, 42). We found that the increase in levels of *treZ* and *osmC* (Fig. 6A and B) or trehalose (Fig. 6C) in ethanol-grown cells did not occur in the Δ*lasR*Δ*rhlR* strain and that the lack of response was not due to ethanol effects on growth in this strain (Fig. S4). Since either *lasR, rhlR* or both AHL responsive transcription factors were necessary for the stimulation of trehalose in cells grown with ethanol, we determined if ethanol enhanced AHL-mediated quorum sensing thereby inducing trehalose levels. To do so, we monitored expression of *rhlI*, a quorum sensing-controlled gene regulated by LasR and RhlR, using a *rhlI*-*lacZ* promoter fusion (98). PA14 wild type and the Δ*lasR*Δ*rhlR* strains were grown ± ethanol. As expected, levels of β-galactosidase activity were much lower in the Δ*lasR*Δ*rhlR* mutant compared to the wild type and we found no significant difference in *rhlI* promoter activity in ethanol-grown cells compared to cells from control conditions (Fig. 6D). The transcriptomics analysis of *P. aeruginosa* grown ± ethanol above did not find evidence for ethanol affecting AHL-mediated quorum sensing broadly (Table S3A and B). Additionally, while AHL quorum sensing was required for trehalose accumulation in response to ethanol, it was not necessary for trehalose accumulation in response to high salt (Fig. 5E); there was, however, a significant reduction in trehalose levels in the Δ*lasR*Δ*rhlR*, compared to wild type in salt suggesting that this mechanism played a role.

**Figure 6.**
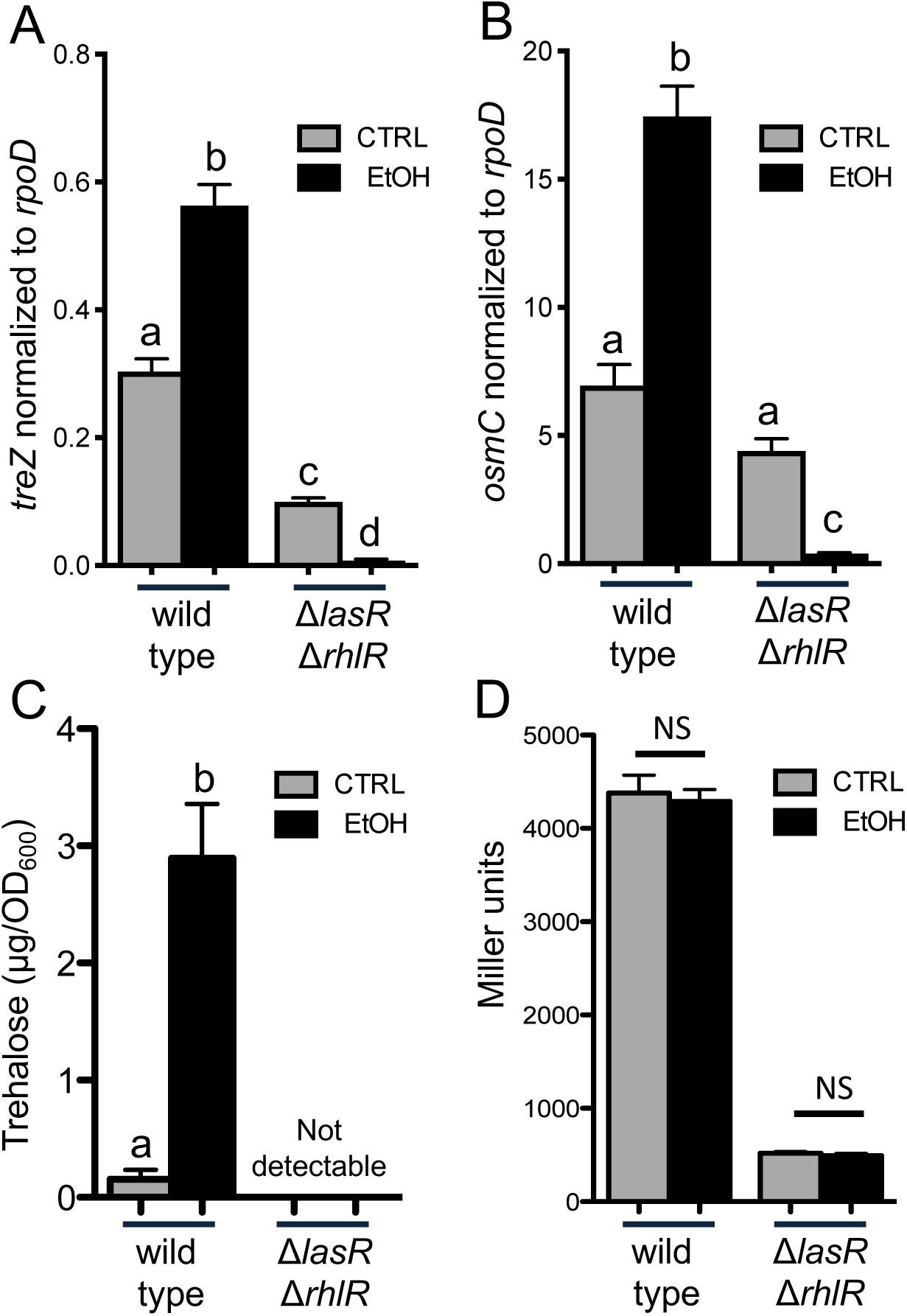
AHL quorum sensing regulation is required for trehalose accumulation and increased *treZ* and *osmC* transcripts in response to ethanol. A. Expression of *treZ* and B. *osmC* genes normalized to *rpoD* in PA14 wild type and Δ*lasR*Δ*rhlR* in buffered LB without and with 1% ethanol for 16 hours. C. Trehalose quantification of PA14 wild type and Δ*lasR*Δ*rhlR* cells grown planktonically in M63 medium without and with 1% ethanol for 16 hours. D. *β*-galactosidase assay of *rhlI*-*lacZ* promoter activity in PA14 wild type and Δ*lasR*Δ*rhlR* cells grown planktonically in M63 medium without and with 1% ethanol for 16 hours. Data are representative of at least 2 independent experiments, each with 3 or more biological replicates. Statistics based on two-way ANOVA and Tukey’s multiple comparisons test. A. a-b, a-d, b-c, b-d p<0.0001; a-c p=0.0004; c-d p=0.0395 B. a-b, b-c p<0.0001; a-c p<0.02. C. a-b p=0.0002.

### Ethanol responsive genes comprise a distinct cluster within a structured network of QS-controlled genes also regulated by AlgU

Together, our data present a complex scheme in which global regulators, AlgU and transcription factors involved in AHL-mediated quorum sensing, control trehalose biosynthetic genes *treYZ* and *osmC* and levels of trehalose in cells grown with ethanol. Our findings support previous reports that separately found *treZ* and *osmC* genes to be among those genes controlled by AlgU (57, 71) and AHL-mediated quorum sensing (41, 42). To test the hypothesis that a subset of genes are members of both the AlgU and QS regulons, and that ethanol specifically altered expression of this subset of genes, we used eADAGE (<u>e</u>nsemble <u>A</u>nalysis using <u>D</u>e-noising <u>A</u>uto-encoders of <u>G</u>ene <u>E</u>xpression), a machine learning algorithm used to generate a model for *P. aeruginosa* expression patterns from 1051 publicly available transcriptome samples (1, 52, 99). The eADAGE model learned 600 expression signatures, and within each signature, genes have different weights. Similarities in weights across signatures for genes indicate a correlation in expression levels. Pairwise Pearson correlation coefficients of the genes in the eADAGE model can be visualized as edge weights in a network where nodes are genes (adage.greenelab.com).

In the network shown in Figure 7, we present the relationships in expression patterns for 1) genes that were found to be differentially expressed in response to ethanol (nodes with orange borders) and 2) the set of genes differentially expressed by more than five-fold in a Δ*lasR*Δ*rhlR* strain compared to the wild type, which included *treZ* and *osmC*, reported in Schuster *et al*. (41) (blue nodes; gene list in Table S3D). Using a dataset that characterized the AlgU regulon by comparing an Δ*algU* mutant to the wild type under AlgU-inducing heat shock conditions (57), we identified genes in the two datasets listed above that were regulated by AlgU (yellow nodes; Table S3C for gene list) or that were altered in bot theΔ*lasR*Δ*rhlR* and Δ*algU* strains (green nodes).

**Figure 7.**
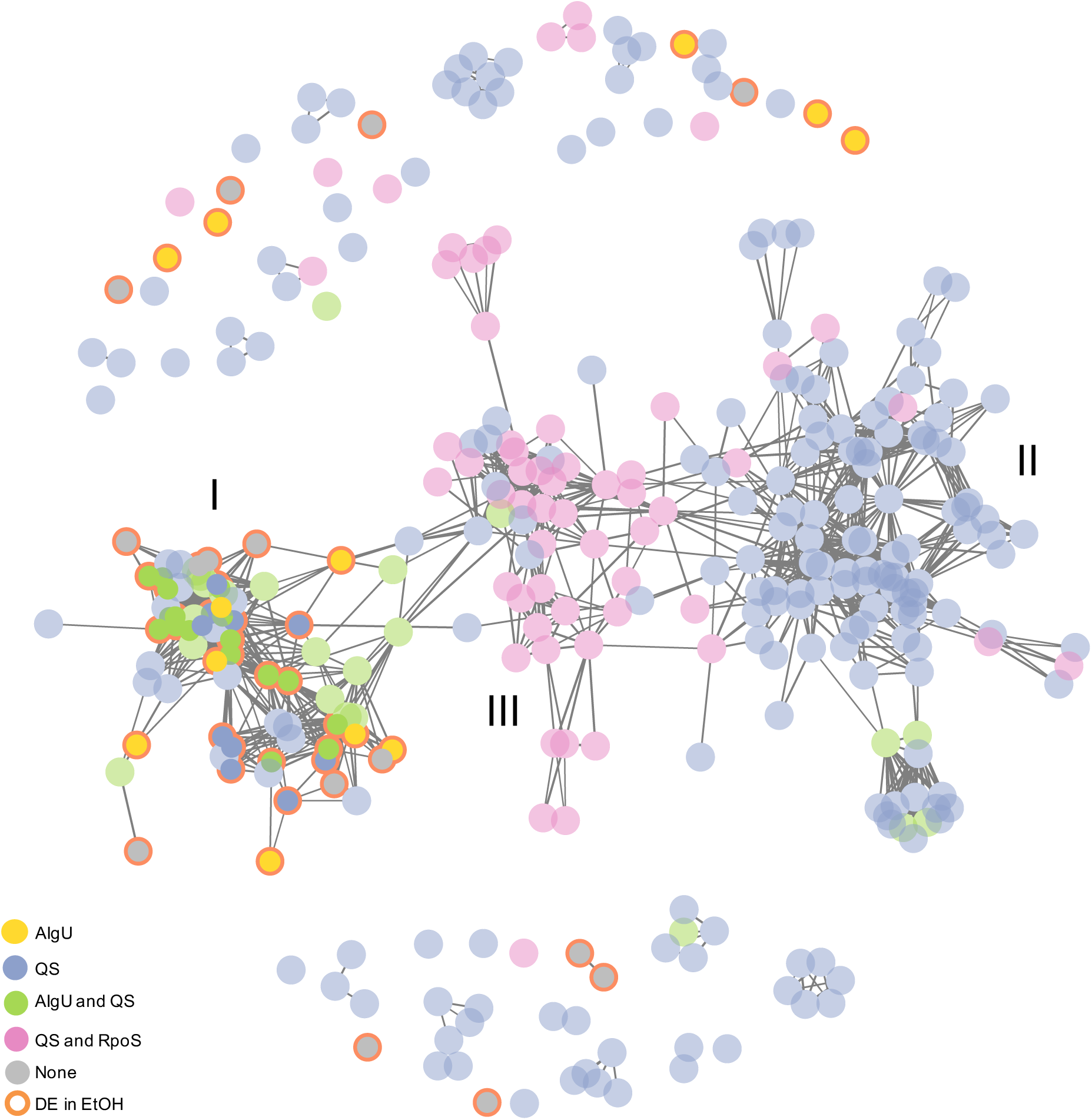
Ethanol responsive genes are enriched for genes with overlapping AlgU and QS control. eADAGE gene-gene network analysis of ethanol responsive genes and QS regulon genes with overlapping AlgU genes indicated. I. Enrichment region for genes coordinately regulated by AlgU and QS (green) and ethanol responsive genes (orange outline). II. Enrichment region for “canonical” quorum sensing genes including *lasI, lasB, rhlR, rhlI*, and *rhlA*. III. Enrichment region for QS genes within the RpoS regulon.

The majority of genes included in this network were connected by edges, revealing strongly correlated expression patterns across the large data compendium comprised of experiments performed by different labs with different strains and in different conditions over more than a decade (1, 52, 99). The connected genes fell into three major clusters (I, II, and III). The majority (80%) of the QS-controlled genes that were also AlgU-controlled (green nodes) (71) were found in cluster I. Ethanol-responsive transcripts (orange borders), including *treZ* and *osmC*, were exclusively localized to cluster I. Statistical analysis found that genes differentially expressed in response to ethanol (Table S3A) represented 12.2% of AlgU-controlled genes (57) and 8.6% of QS-controlled gene sets defined above (Table S3 for gene lists), but comprised 44% of genes present in both regulons. The enrichment of the intersection of AlgU and QS-controlled genes over the sets of either all AlgU-or all QS-regulated genes was significant (p=0.001, and p=0.00002, respectively). Visualization of a cluster of ethanol-responsive genes within the gene-gene network comprised of the complete AlgU regulon (57) is shown in Figure S5.

Cluster II genes contained many genes known to be regulated mainly by LasR or RhlR, and their cognate signals (41, 42). Examples of genes in cluster II are *lasI, lasB, rhlR, rhlI*, and *rhlA*. The lack of any of the ethanol-responsive genes in cluster II is consistent with our findings that ethanol did not alter expression of *rhlI* (Fig. 6D) and is in support of our model that ethanol did not broadly induce the entire QS regulon.

Cluster III contained genes from the QS-controlled gene set that were previously described by Schuster (42) as also being differentially expressed in an *rpoS* mutant (pink nodes) (42) (Fig. 7 and Table S3E for gene list). Of the 56 genes in cluster III, 47 (42) and 29 (57) genes were differentially expressed in separate published studies describing the RpoS regulon. Ethanol-responsive genes were not among genes in cluster III, supporting above data showing that ethanol-induced trehalose levels were not dependent on *rpoS* (Fig. S3). Specific enrichment in AlgU and QS co-regulated genes among the genes upregulated in response to ethanol is consistent with ethanol activating only a subset of the AlgU and AHL-controlled regulons.

### Ethanol induces trehalose after entry into post-exponential phase

Increased trehalose levels and *osmC* and *treZ* transcripts in cultures with ethanol was dependent on both SpoT-synthesized (p)ppGpp and AHL-mediated quorum sensing, signals associated with growth restriction, often due to nutrient limitation, and high cell density, respectively. Analysis of trehalose levels in control and ethanol-containing cultures found that ethanol only affected trehalose levels in cells from post-exponential phase cultures, but not exponential phase cultures. This relationship was observed in both M63 medium (Fig. 8A) and buffered LB medium (1.96 μg/OD_600_ in control cultures vs 2.81 μg/OD_600_ in ethanol-containing cultures, p-value=0.1339). Expression of *treZ* was also not affected by ethanol in cells from exponential phase cultures, but was in cells from the same cultures collected after entry into post-exponential phase (Fig. 8B). Together, these data support a model in which the induction of trehalose in response to ethanol by AlgU requires other signals from quorum sensing and growth-restriction associated pathways.

**Figure 8.**
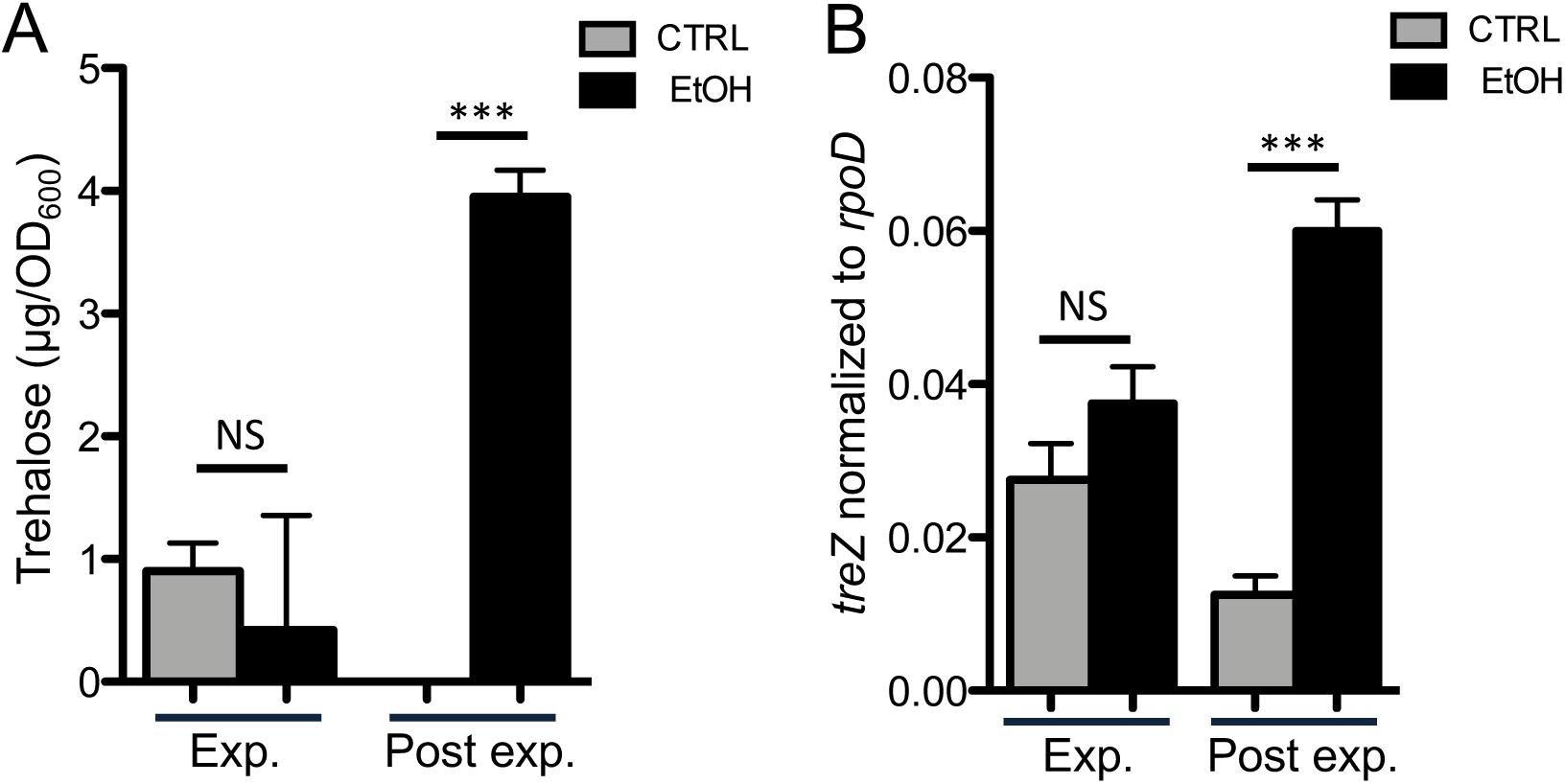
Ethanol stimulates *treZ* gene expression and trehalose levels in cells from post-exponential phase but not exponential cultures. A. Trehalose quantification of cells grown in M63 medium without and with 1% ethanol for 6 hours to an OD_600_ of ∼1.0 (Exp.) or 16 hours (Post exp.). B. *treZ* gene expression normalized to *rpoD* in cultures grown in buffered LB without and with 1% ethanol to the same densities as in A. Data are representative of 3 experiments, each with 2-4 biological replicates. Statistics based on two-way ANOVA and Tukey’s multiple comparisons test. ***p≤0.0008.

## Discussion

The data presented above lead us to propose a model, based largely on genetic analyses, in which ethanol activates AlgU through stimulation of (p)ppGpp, synthesized by SpoT, and activation of DksA-dependent transcription. Chromatin immunoprecipitation experiments have shown that AlgU binds to the promoter for the *treYZ* containing operon (57). AHL-mediated quorum sensing, through LasR and/or RhlR was required for AlgU-dependent activation of *treZ* and *osmC* and increased levels of trehalose, and thus ethanol induction of trehalose was only observed in cultures after AHL-mediated QS was induced. Our data do not indicate that ethanol led to a global increase in expression of the QS regulon. Based on these data, we propose that even though ethanol was present over the course of growth, the increased trehalose biosynthesis in response to ethanol only occurs when cells have sensed a quorum and when (p)ppGpp synthesis can be stimulated, perhaps because of concomitant nutrient limitation signals which are known to activate SpoT (Fig. 9). These data highlight a nuanced response to a microbially-produced molecule, ethanol, in *P. aerugionsa* and this underscores how microbe-microbe interactions may change with shifts physiological states and extracellular signal concentrations.

**Figure 9.**
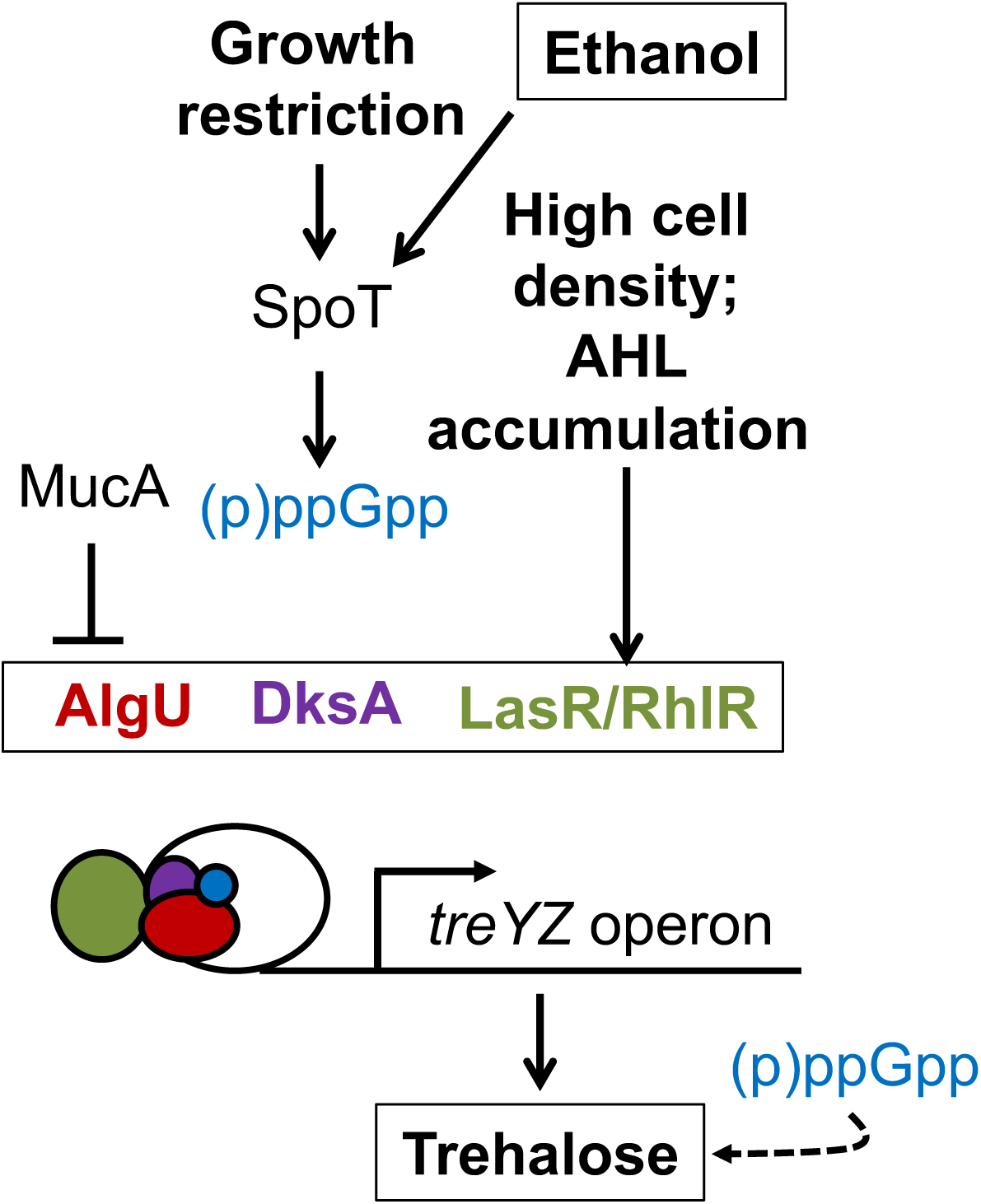
Model describing ethanol stimulation of the AlgU regulon and trehalose production. Our data support a model wherein cell density and growth signals determine the cell’s response to ethanol. The colors of the text are reflected in the schematic of the transcriptional regulatory complex.

The model for AlgU activation by (p)ppGpp and DksA represents a new mechanism by which the important sigma factor, AlgU, may activated in *P. aeruginosa*. (p)ppGpp-dependent activation of E. coli RpoE, an AlgU homolog, has been reported previously (35) and has been associated with growth phase (100), but activation by non-inhibitory concentrations of ethanol has not been reported. Our data showed that ethanol, at a 1% concentration, did not activate AlgU through effects on MucA or MucB proteins (Fig. 3), which are targeted for cleavage in response to periplasmic or envelope stress. In the presence of 0.5 M salt, however, a stimulus that is expected to induce periplasmic stress, DksA-and SpoT-independent stimulation of trehalose is observed, presumably because some AlgU activation occurred through MucA cleavage, but DksA and SpoT did contribute to the strength of the response suggesting that these mechanisms can work together.

The intersection of QS regulation and AlgU regulation is interesting. Future experiments will also determine if LasR and/or RhlR directly interact with the promoter upstream of the *treYZ* containing operon, and if that interaction only occurs when AlgU and DksA are complexed with RNA polymerase. Evidence for direct activation of *osmC* and *treYZ* expression is that overexpression of *lasR*, but not *rhlR*, is sufficient to induce expression of these genes (42). Our variable results with single mutants lacking either *lasR* or *rhlR* (data not shown) led us to propose that both transcription factors can influence the expression of *treYZ*, but with different kinetics, as has been shown for other genes (101). In separate studies with Δ*dksA* and Δ*relA*Δ*spoT* mutants in strain PAO1, DksA and (p)ppGpp have independently been associated with both the positive and negative regulation of genes that are differentially expressed in a Δ*lasR*Δ*rhlR* mutant (96, 102-106) so there may be a complex relationship between these signaling pathways.

The biological role of AlgU-induced trehalose in *P. aeruginosa* in response to stresses or in microbial communities is not yet known. AlgU has been implicated in both the positive and negative regulation of genes involved in oxidative and osmotic stress responses in other Pseudomonads (107-110). For example, in different pathovars of *P. syringae*, AlgU regulates oxidative and osmotic stress response genes transcriptionally in response to osmotic stress (109) and contributes to plant disease independently of alginate (110). In *Pseudomonas fluorescens*, an *algU* mutant was significantly more sensitive to osmotic stress than wild type (107). *P. fluorescens* AlgU to be necessary for desiccation stress tolerance, but dispensable for tolerance to 3% hydrogen peroxide, 1.9% paraquat, 5% sodium hypochlorite, heat shock, pH extremes, and the reducing agent dithiothreitol (107). In *P. aeruginosa*, AlgU has been described as having a negative role in oxidative stress resistance, can be protective against host innate immune factors through its regulation of alginate (25, 26). We did not observe a protective benefit of growth with 1% ethanol in oxidative stress, osmotic stress, and dessication assays in the growth conditions used in these studies (data not shown). Trehalose has been reported to protect against osmotic, oxidative, heat, and cold stress by stabilizing proteins and reducing formation of denatured protein aggregates (111-115), as a carbon reserve (111, 116), and recent studies have found that trehalose can stabilize outer membrane vesicles (117). Trehalose can accumulate in the cytoplasm and periplasm and be secreted, making it an interesting molecule to consider in the context of microbe-microbe interactions. *P. syringae* survival as an epiphyte and *P. aeruginosa* plant pathogenesis both require the ability to make trehalose (64, 118). Interestingly, exogenous trehalose and wild type-derived trehalose in co-culture rescued the attenuated trehalose mutant phenotype in *P. aeruginosa Arabidopsis* pathogenicity *in planta*, with the requirement for trehalose being independent of osmoprotection (64). Pseudomonads can accumulate a variety of osmoprotectants in addition to trehalose, including betaine, ectoine, and *N*-acetylglutaminylglutamine amide (NAGGN) (118). This redundancy may contribute to the fact that trehalose mutants are not more sensitive to tested stresses in laboratory assays (64).

In addition to *treZ* and other trehalose metabolism genes, cluster I included 38 genes found within the genome island that spans from PA2134 to PA2192 (PA14_36980-PA14_36345) (73, 119, 120). 16 of the 54 genes (∼30%) differentially expressed in response to ethanol were in this chromosomal region. Many of the genes in this chromosomal region that are differentially expressed in ethanol could participate in survival of stresses likely to be present in mixed-species communities formed with ethanol-producing microbes. For example, *glgE* is involved in the metabolism of glycogen, a carbon and energy storage molecule that accumulates when carbon is in excess relative to other growth-limiting nutrients (121). Other genes within this genome island encode putative ion transporters, double strand break repair enzymes, and two catalases. Other ethanol-induced genes within Cluster I included *osmC, sprP*, and *pfpI*. OsmC can be protective against oxidative stress caused by exposure to elevated osmolarity and hyperoxides through an unknown mechanism (122). SprP is a subtilase protease (123) and *pfpI* codes for a protease that plays a role in DNA protection in non-stress conditions and in the presence of hydrogen peroxide (124). Mutation of either *sprR* or *pfpI* in *P. aeruginosa* has pleiotropic effects (123, 124).

The important next question is the mechanism by which ethanol stimulates (p)ppGpp. Others have shown that (p)ppGpp levels increase in response to higher concentrations of ethanol, and, in *E. coli*, the addition of ethanol mimics amino acid starvation (125). They speculated that ethanol (and other short-chain alcohols) may interfere with amino acid uptake (125). Ethanol may also directly impact ribosome activity (126) or other pathways through effects on cell membranes, and the fact that responses vary based on growth phase provides a useful tool to understand how ethanol and ethanol-producing microbes influence the other bacteria.

## Acknowledgements

Research reported in this publication was supported by National Institutes of Health (NIH) grant R01 GM108492 to D.A.H., NIAID T32AI007519 to C.E.H. and D.L.M, NIGMS T32-GM008704-16 to G.D., NHLBI T32HL134598-01 to M.E.C., Burroughs Wellcome Fund award to D.N., Canadian Institutes of Health Research fellowship to D.M.. Microarray processing was carried out at Dartmouth Medical School in the Genomics Shared Resource Core, which was established by equipment grants from the NIH and NSF and is supported in part by a Cancer Center Core Grant (P30CA023108) from the National Cancer Institute. Support for the project was also provided by the NIGMS P20GM113132 through the Molecular Interactions and Imaging Core (MIIC). The content of this publication is solely the responsibility of the authors and does not necessarily represent the official views of the NIH.

We thank Carol Ringelberg (Geisel School of Medicine at Dartmouth) for Microarray data processing work. We also thank Fred Ausubel (Harvard Medical School), Daniel Wozniak (The Ohio State University College of Medicine), and Dennis Ohman (Virginia Commonwealth University School of Medicine) for generously providing strains. We also acknowledge Gary Heussler (UCSD Division of Biological Sciences) for providing plasmid and Jong-In Hong (Seoul National University) for providing the PyDPA probe.

